# IsoMobil: Resolving Molecular Ambiguity in Mass Spectrometry-based Spatial Omics Through Ion Mobility

**DOI:** 10.64898/2026.08.07.743457

**Authors:** Meenakshi, Lukasz G. Migas, Kameron R. Molloy, Katerina V. Djambazova, Jeffrey M. Spraggins, Raf Van de Plas

**Affiliations:** Delft Center for Systems and Control, Delft University of Technology, Delft, Netherlands; Mass Spectrometry Research Center, Vanderbilt University, Nashville TN, USA; Department of Cell and Developmental Biology, Vanderbilt University, Nashville TN, USA; Department of Biochemistry, Vanderbilt University, Nashville TN, USA; Department of Chemistry, Vanderbilt University, Nashville TN, USA; Department of Pathology, Microbiology, and Immunology, Vanderbilt University Medical Center, Nashville TN, USA

**Keywords:** imaging mass spectrometry, ion mobility, spatial omics, spectral pattern recognition, isotopologue, dimensionality reduction

## Abstract

Molecular imaging by imaging mass spectrometry (IMS) has become a key modality for spatial proteomics, lipidomics, glycomics, and metabolomics. It maps hundreds to thousands of molecular species concurrently throughout tissue without prior labeling. However, reporting thousands of ion images makes IMS measurements very high-dimensional, complicating interpretation. Furthermore, IMS data contain implicit chemical relationships. For example, the same molecular species can be reported by several separately-measured ion species, each an isotopic variant or isotopologue of that molecule. While conventional dimensionality reduction methods such as principal component analysis can address the dimensionality challenge, they typically do not preserve chemical relationships (*e*.*g*., isotopologue grouping), making biological interpretation harder. As advanced, higher-dimensional measurement types such as ion mobility IMS (IM-IMS) expand into spatial omics, addressing interpretability in a chemically informed way becomes pressing. Therefore, we present IsoMobil, a dimensionality-reduction framework for IM-IMS data that empirically detects potential isotopologues. Besides reducing dataset complexity, it facilitates interpretation at the (biologically relevant) molecular-species level rather than ion-species level. The algorithm finds spatially coherent ion species, filters them based on isotope-induced mass-to-charge (*m/z*) distances and mobility-bin consistency (isotopologues have near-identical collisional cross-sections). This yields a compact representation where isotopologue-candidate families, rather than individual ion-species, form latent dimensions. In a synthetic benchmark, IsoMobil outperformed (F1=1.0) spatial-only and *m/z*-based methods (F1≈0.67). In a human colon case study, IsoMobil found 77 isotopologue-candidate groups (COSH-P-quality≥0.85) among 6344 lipid ion species. By automating isotopologue discovery, IsoMobil lifts biological interpretation of exploratory, untargeted spatial omics by IM-IMS to the molecular-species level.

## 1 Introduction

Imaging mass spectrometry (IMS) is a molecular imaging modality that, in a single experiment, concurrently maps hundreds to thousands of molecular species across a tissue section [4, 11, 25], linking molecular distributions to tissue morphology. Since IMS is highly multiplexed and does not require prior labeling, it is well-suited for exploratory spatial biology studies and has become a key modality for spatial proteomics [21], lipidomics [31], glycomics [2], and metabolomics [3]. However, since IMS measurements report thousands of ion species’ distribution images, they are very high-dimensional and complex, hindering interpretation. Furthermore, IMS data contain implicit chemical relationships. For example, the same molecular species can be reported by several separately-measured ion species, each an isotopic variant or isotopologue of that molecule [16, 29]. Isotopes are atoms of the same element with different neutron numbers and, thus, different mass, such as ^12^C and ^13^C, and isotopologues are molecules that differ only in isotopic composition. In measurements of primarily singly-charged carbon-dominated ions, isotopologues typically appear as a series of peaks with near-1 *m/z*-spacing. The dimensionality challenge in IMS is conventionally addressed by dimensionality reduction methods such as principal component analysis (PCA) [19, 20, 18], non-negative matrix factorization (NMF) [30, 9, 7, 17], t-distributed stochastic neighbor embedding [10, 1], and uniform manifold approximation and projection [24, 12]. However, these approaches typically do not preserve chemical relationships (*e*.*g*., their lower-dimensional representations do not necessarily group isotopologues of the same molecular species together), complicating biological insight. As more advanced measurement types, such as ion mobility IMS (IM-IMS) [26], enter spatial omics, adding a further mobility dimension to IMS, it is essential to develop methods that digest such data and reduce their dimensionality in a chemically informed way such that they help rather than hurt interpretation.

To address this gap, we introduce IsoMobil, a dimensionality reduction-framework that casts IM-IMS data to a lower-dimensional representation that only permits isotopologue-consistent feature groupings. This implicitly automates isotopologue discovery in IM-IMS omics-data. Besides reducing dataset complexity, IsoMobil effectively facilitates interpretation at the (biologically relevant) molecular-species level rather than the (natively-measured) ion-species level. Conventional isotopologue discovery/removal methods [29, 23, 22, 28, 27] are often unaware of spatial relationships, rely on theoretical isotope-intensity envelopes, or require prior knowledge on molecular formulas, charge states, or compound classes present in the sample. These assumptions are rarely met in a molecularly heterogeneous tissue environment. More recent approaches such as DeepIon [6] and IsoSpace [13] exploit spatial relationships, but combining spatial and *m/z*-based criteria still yields substantial isotopologue false positives, *e*.*g*., due to isomers and isobars. Isomers are molecular species that have the same molecular formula, and thus the same exact mass, but a different arrangement of their atoms. Isobars are molecular species that have different molecular formulas, yet a very similar (but not the same) exact mass. In the presence of isomers and isobars, several ion species could be ‘hiding’ behind, respectively, the same or nearly the same *m/z*-value. If only *m/z*-information is used, both isomers and isobars can confound an isotopologue grouping. IsoMobil addresses this ambiguity by combining spatial, mass spectral (*m/z*), and mobility criteria, effectively enabling automatic annotation (without prior knowledge) of spatial-, *m/z*-, and mobility-consistent isotopologue candidates. This paper introduces IsoMobil with demonstrations on three datasets. A synthetic case study with ground truth compares IsoMobil’s performance to classical dimensionality reduction and mobility-unaware isotopologue discovery methods. A mouse pup IM-IMS case study demonstrates the impact of mobility information on isotopologue specificity. Finally, a human colon IM-IMS case study shows how IsoMobil’s isotopologue discovery provides chemically relevant dimensionality reduction and facilitates molecular-species level interpretation of spatial lipidomics data.

## 2 IsoMobil Method

Like most dimensionality reduction methods, IsoMobil groups natively measured features (ion species in IM-IMS data) together into a smaller set of latent features. Unlike traditional methods, it only permits feature groupings where the grouped ion species are potential isotopologues of the same molecular species. To accomplish this, IsoMobil imposes on its latent components’ spatial, mass spectral, and ion mobility signatures criteria consistent with those of potential isotopologues. Each IM-IMS-experiment yields a 3-mode tensor of nonnegative intensity values, with a spatial mode of size *N* (number of pixels), a mass spectral mode of size *S*_1_ (number of *m/z*-bins), and a mobility mode of size *S*_2_ (number of mobility-bins, *mb*) (Figure 1). After pre-processing, peak-picking in the (*m/z, mb*)-domain yields a matrix **X** ∈ ℝ^*N* ×*M*^, with *N* the number of pixels and *M* the number of found ion species. Each ion species feature has both an *m/z*- and *mb*-value associated with it, separately accessible through *m/z*-bin vector **y** ∈ ℝ^*M*^ and mobility-bin vector **z** ∈ ℝ^*M*^. The goal is to find groups of ion species whose members exhibit a coherent (co-localized) spatial distribution, an *m/z*-spacing consistent with isotopic substitution, and an *mb*-distance small enough to suggest identical molecular shape, all criteria expected for isotopologues reporting the same molecular species. The algorithm has three stages. **Stage I** enforces the spatial coherence criterion by applying NMF to matrix **X**. This yields *k* non-negative components, each grouping ion species that are spatially coherent, but that do not necessarily meet the *m/z*- and *mb*-criteria. Within each component *r*, those ion species with NMF coefficients above a user-defined component intensity threshold *ρ* are considered for filtering in Stage II. **Stage II(a)** enforces the *m/z*-criterion using a binary candidate pattern matrix **Π** ∈ ℝ^×*M*^. Based on **y, Π** encodes a pairwise link between ion species *i* and *j* if their *m/z* separation falls within a user-defined tolerance around the *m/z*-spacing expected for singly-charged ^13^C-dominated isotopologue peaks (nominally 1 Da, exactly 1.003355 Da): ||*m*_*j*_ − *m*_*i*_| − 1| ≤ *ϵ* with *m*_*i*_ and *m*_*j*_ their *m/z*-values and *ϵ*, a user-defined tolerance. The *ϵ*-tolerance is meant to absorb any *m/z*-measurement and peak-picking uncertainty, as well as the small mass defect of ^13^C spacing. This work measures primarily singly-charged, carbon-rich ions, so charge-state-dependent spacings or non-carbon isotope envelopes are not considered here. Missing intermediate isotopologues are allowed and handled as in [13]. **Stage II(b)** filters the links in **Π** further for *mb*-consistency. Using **z**, a candidate link is retained only if |*mb*_*j*_ − *mb*_*i*_| ≤ *δ*, where *δ* is a user-defined *mb*-tolerance. Finally, Algorithm 1 exhaustively extracts isotopologue candidate groupings from **Π**, storing each “isotopologue family” as a separate row in results matrix 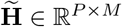, with *P* the total number of extracted isotopologue candidate groupings.

**Fig. 1.**
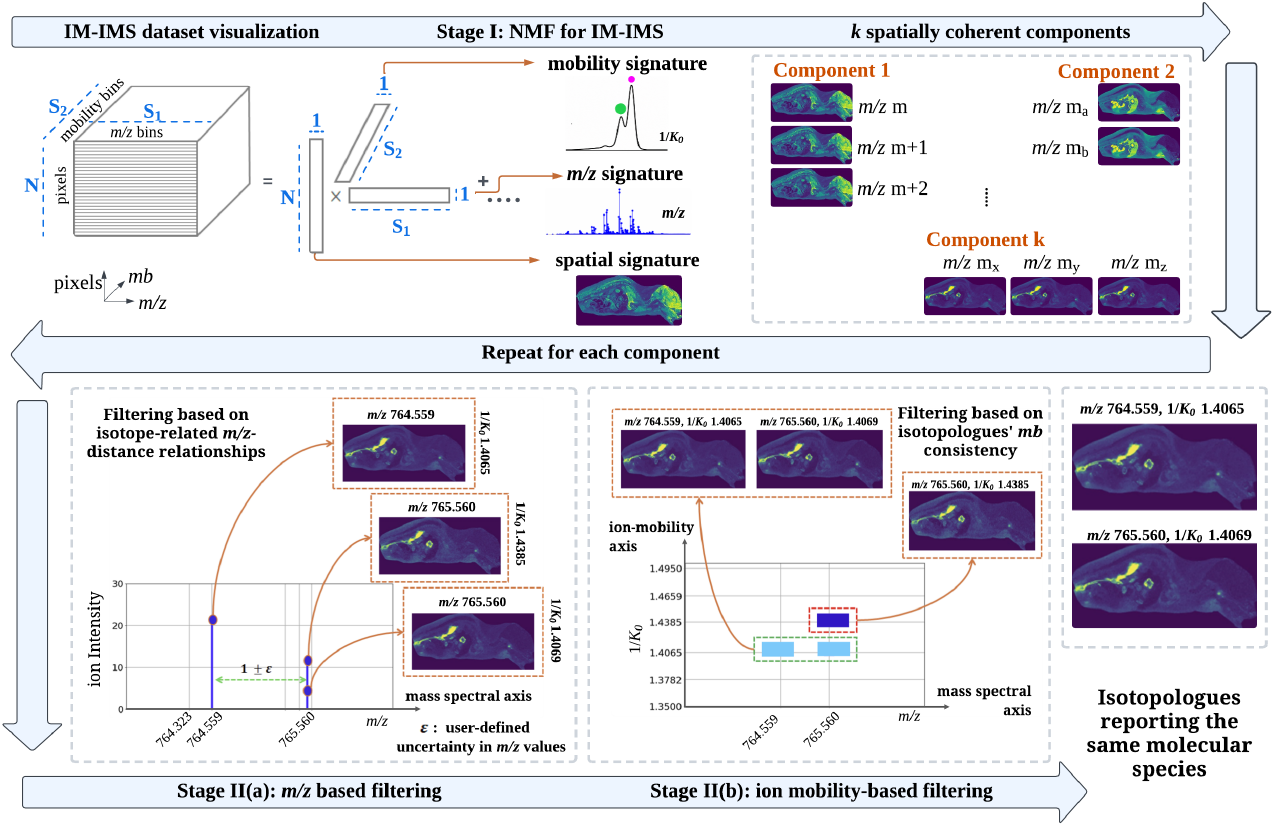
IsoMobil pipeline. **Stage I** identifies spatially co-localized ion species by NMF. **Stage II(a)** filters by carbon isotopes’ 1.003355 Da-distance, using ||*m*_*j*_ *− m*_*i*_| *−* 1| ≤ *ϵ*, with tolerance *ϵ*. **Stage II(b)** retains only ion species with matching mobility, using |*mb*_*j*_ *− mb*_*i*_| *≤ δ*, with tolerance *δ*. Each family of isotopologue candidates is a row in results matrix 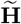 and maps several ion species features to one molecular species feature.

## 3 Experiments and Results

### 3.1 Datasets & Evaluation Metrics

#### Synthetic dataset (case study 1)

As we have no ground truth for the isotopologue content of empirically measured data, we first generate a synthetic IM-IMS dataset with known content. It consists of 80 × 25 pixels with five non-overlapping Gaussian spatial distributions, mimicking localized tissue regions. We introduced eight isotopologue families to recover, with 2–3 peaks separated by *m/z* ≈ 1.00335. The peak intensities follow a binomial ^13^C natural-abundance model [28, 14], with carbon counts sampled from [34, 48] to reflect the phospholipid mass range observed in colon tissue. To mimic realistic acquisition conditions, we add 8-bin intra-family *mb*-jitter, 80 unstructured background peaks, and 5% Poisson shot noise. **Mouse pup dataset (case study 2)** is the IM-IMS measurement of a whole-body mouse pup section (*m/z* 200-1500, 50 *µ*m pixel size, *mb* 305.5-5366.5, 1*/K*_0_ 1.37-1.52 Vs/cm^2^) [5]. After TIC-normalization and peak-picking, it reports 2413 ion species features. **Human colon dataset (case study 3)** is an IM-IMS measurement of de-identified human colon tissue (*m/z* 400–1300, 10 *µ*m pixel size, *mb* 370.5-6921.5, 1*/K*_0_ 0.8-1.8 Vs/cm^2^) [15]. Its 6344 ion species features demonstrate the complexity of IM-IMS-based spatial lipidomics datasets. The quality of a retrieved isotopologue candidate family 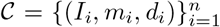, with *n* ion species members and *I*_*i*_, *m*_*i*_,. and *d*_*i*_ respectively the ion image, *m/z*-value, and *mb*-value of the *i*th member, is evaluated using three metrics. Groups with *n*=1, ‘singletons’, are not considered. **COSH-P** measures spatial similarity among members by averaging the cosine similarity between Histogram of Oriented Gradients (HOG) descriptors, **h**_*i*_ = HOG(*I*_*i*_), of the member images: 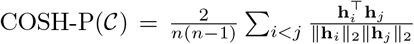 Higher COSH-P values indicate greater spatial similarity among isotopologue candidates. **Intra-pattern spatial correlation (IPC)** [13] measures the average Pearson correlation between vectorized member ion images, **x**_*i*_ = vec(*I*_*i*_): 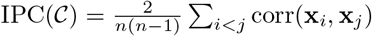. Higher IPC indicates stronger co-localization among members. **Intra-pattern mobility variability (IPMV)** measures mobility variability within a candidate family: IPMV(*C*) = max_*i*_ − *d*_*i*_ min_*i*_ *d*_*i*_. Lower IPMV means stronger mobility consistency. While not a definitive isotopologue assessment, these metrics report a family’s spatial and mobility coherence and can help rule out false positives.

##### Algorithm 1

IsoMobil’s algorithm to extract isotopologue candidate families.

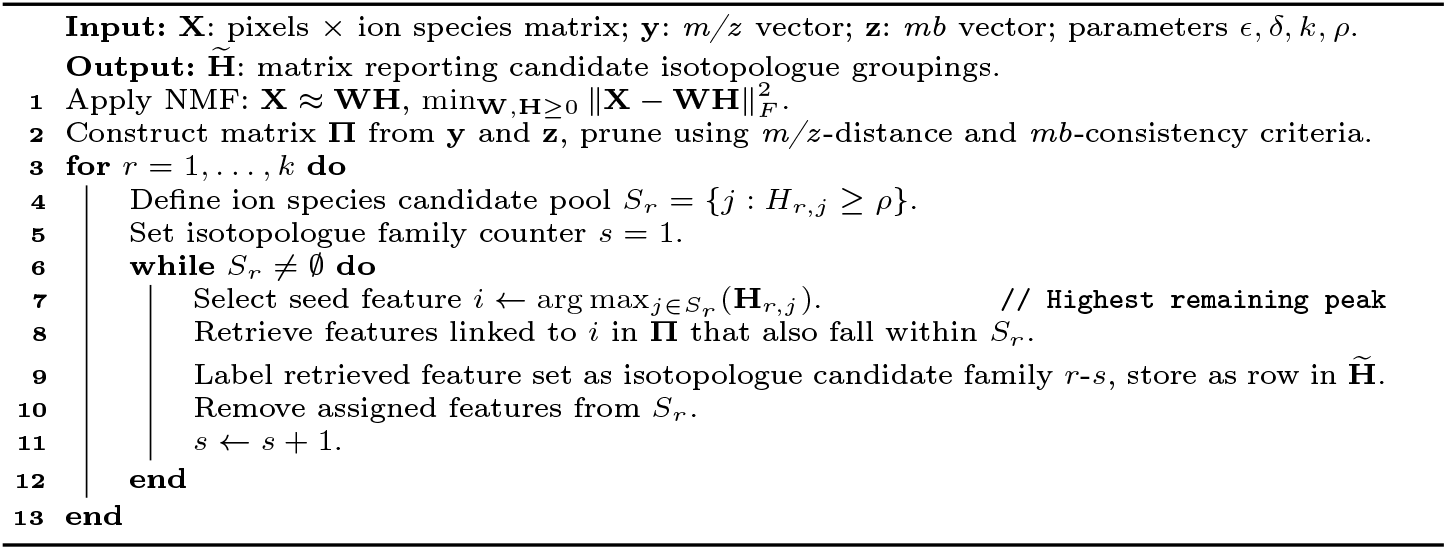

### 3.2 Results and Discussion

#### Case Study 1 – synthetic benchmark

IsoMobil is evaluated on a synthetic dataset with known isotopologue content under two conditions: *Clean* contains only true isotopologue families and background peaks, and *Confounded* adds one decoy pair per family. Each decoy satisfies the spatial co-localization and near-1 *m/z*-spacing criteria, mimicking isobaric lipid interference, but is separated by 100 *mb*, violating the mobility consistency of true isotopologues. Parameters are set to *k* = 5, *ϵ* = 0.020, *δ* = 30 and detections are matched to the ground truth using Intersection over Union ≥0.50. Figure 2 shows that in the *Clean* condition, IsoSpace, IsoMobil-PCA (PCA instead of NMF), and IsoMobil achieve a perfect F1 = 1.0 score, while unconstrained PCA and NMF perform poorly for isotopologue detection. Under *Confounded* conditions, IsoMobil maintains F1 = 1.0 and Precision = 1.0, whereas *m/z*-only IsoSpace drops to F1 ≈ 0.67. This demonstrates that the Stage II(b) *mb*-filter is critical for rejecting mobility-incoherent decoys and, thus, for isotopologue specificity.

**Fig. 2.**
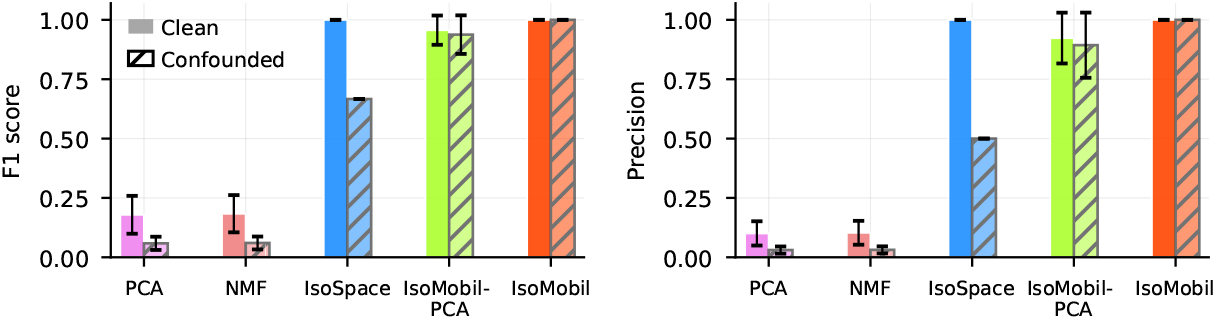
Case study 1 results - mobility constraint preserves isotopologue specificity against confounders. Isotopologue discovery trials by PCA, NMF, IsoSpace, IsoMobil-PCA, and IsoMobil are compared by F1 score (left) and precision (right) under *Clean* (solid) and *Confounded* (hatched) conditions. Spatial and *m/z*-only methods’ performances degrade when mobility-separated decoys (*e*.*g*., isobars, isomers) are present. IsoMobil maintains its performance. (Bars represent mean *±* st. dev.)

#### Case Study 2 - presence of mobility information enables less stringent mass resolution requirements

This case study assesses the impact of mobility information on isotopologue detection. We compare IsoSpace [13], which uses spatial and *m/z*-filtering, to IsoMobil, which adds mobility-filtering. Figure 3 shows an example from the mouse pup dataset. IsoSpace groups ion species that are spatially similar and at the right *m/z*-distances. However, their *mb*-values reveal that they do not stem from the same molecular species. Their 504.5 *mb*-gap (*e*.*g*., *m/z* 766.571 at *mb* 3239.5 and *m/z* 767.574 at *mb* 3744.0) suggests distinct 1*/K*_0_ and species, which shows isobaric species confounding detection, while IsoMobil’s mb-filter refines isotopologue discovery. Figure 3 (right) shows the number of high-quality candidate families (COSH-P>0.70) retrieved from the mouse pup data with different *ϵ*-tolerances (IsoSpace & IsoMobil) and *δ*-tolerances (IsoMobil only). Since IsoSpace relies solely on *m/z*-filtering, its *m/z*-tolerance needs to be narrow (*ϵ* = 0.007) to yield high-quality results. By adding *mb*-filtering, IsoMobil is not only able to achieve high-quality results with a less stringent *m/z*-tolerance (*ϵ* = 0.009), but it also yields higher numbers of retrieved families. The presence of mobility information enables better isotopologue discovery with less stringent mass resolution requirements, *i*.*e*., isotopologue specificity no longer relies on the *m/z*-axis alone.

**Fig. 3.**
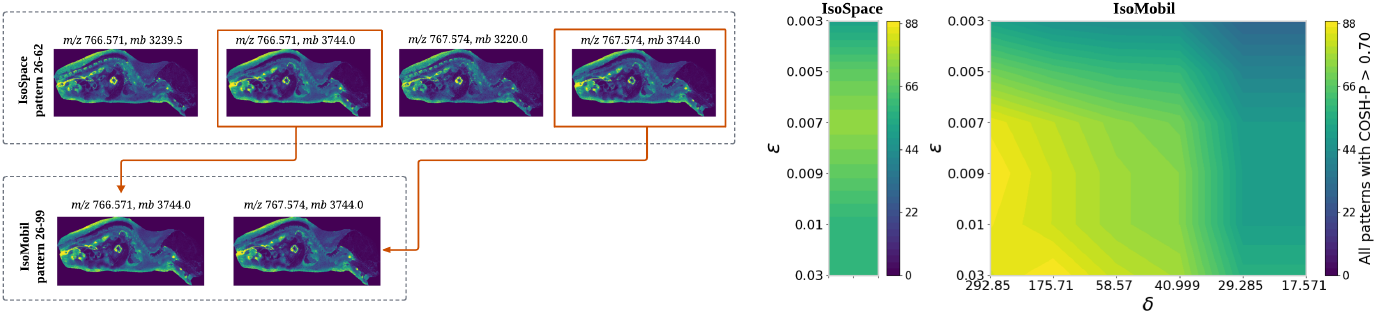
Case study 2 results-presence of mobility information enables less stringent mass resolution requirements. (left) IsoSpace groups co-localized, near-1 *m/z*-spaced ion species (note their *mb*-differences) together into a *mb*-inconsistent candidate family. IsoMobil’s *mb*-filtering retains only mobility-coherent candidates. (right) The use of mobility improves candidate discovery across different *ϵ* and *δ* settings, while also allowing *m/z*-filtering to be less strict.

#### Case Study 3 - performance on human colon tissue

This study demonstrates IsoMobil’s performance on human colon tissue. Figure 4a shows the quality and number of found (non-singleton) isotopologue candidate families as a function of *m/z*-tolerance *ϵ*. The number of found families peaks at *ϵ* = 0.009. Wider *m/z*-tolerances allow less coherent patterns and noise to sneak in. Narrower *m/z*-tolerances can be overly restrictive and may reject genuine isotopologues, highlighting a critical trade-off. Figure 4b shows sensitivity to *mb*-tolerance *δ*, varied from 0.1% to 2% of the total 6921.5 *mb*-range. Better COSH-P scores were obtained with *δ* ranging from 34.608 to 69.215. Again, there is a trade-off: narrow *mb*-tolerances may reject plausible candidates, whereas broad tolerances may group together ion species stemming from distinct molecular species. For example, using *δ* = 34.608 retrieves 77 isotopologue candidate families of high quality (COSH-P*>* 0.85), while *δ* = 48.450 finds more families, albeit of lesser quality (COSH-P*>* 0.60). Figure 4c compares IsoMobil to PCA, NMF, and IsoSpace using *ρ* = 0.10, *k* = 30, *ϵ* = 0.009, and *δ* = 34.608 (0.5% of *mb*-range). PCA and NMF yield ~ 2200 components that group co-localizing ion species, but their IPMVs span hundreds of *mb*, suggesting different molecular species being thrown together. By adding an *m/z*-filter, IsoSpace reduces this to 101 isotopologue candidate families. IsoMobil’s *mb*-filter refines this further to 69 families with near-zero mean IPMV (≈ 7 bins), a 14-fold reduction from IsoSpace. Mean IPC is comparable for IsoMobil and IsoSpace (0.69), confirming that IsoMobil reduces IPMV without sacrificing spatial coherence. Figure 5 shows an example where spatial+*m/z* filtering is insufficient. Two ion species with nearly identical *m/z*-values (both rounding to *m/z* 854.697) but different mobilities (*mb* 1620.0 and 1702.5), *i*.*e*., unresolved co-detected species, meet the spatial and *m/z*-criteria. While slight differences in spatial distribution (arrows) subtly suggest that one might be a false positive member of this pattern, removal based on spatial cues can be difficult and only works when localizations are different. Instead, ion species (*m/z* 854.697, *mb* 1702.5) is easily removed by narrowing the *mb*-tolerance, demonstrating that *mb*-filtering resolves ambiguities that are hard to address by spatial+*m/z*-filtering alone. Finally, biological relevance was supported by orthogonal LC-MS/MS evidence of IsoMobil’s human colon isotopologue groupings. These include successfully finding isotopologues of species such as sphingomyelin SM(d18:1_16:0) (family at *m/z* 725.558) [8] and phosphatidylcholines PC(34:1) (family at *m/z* 782.565) and PC(36:4) (family at *m/z* 804.553).

**Fig. 4.**
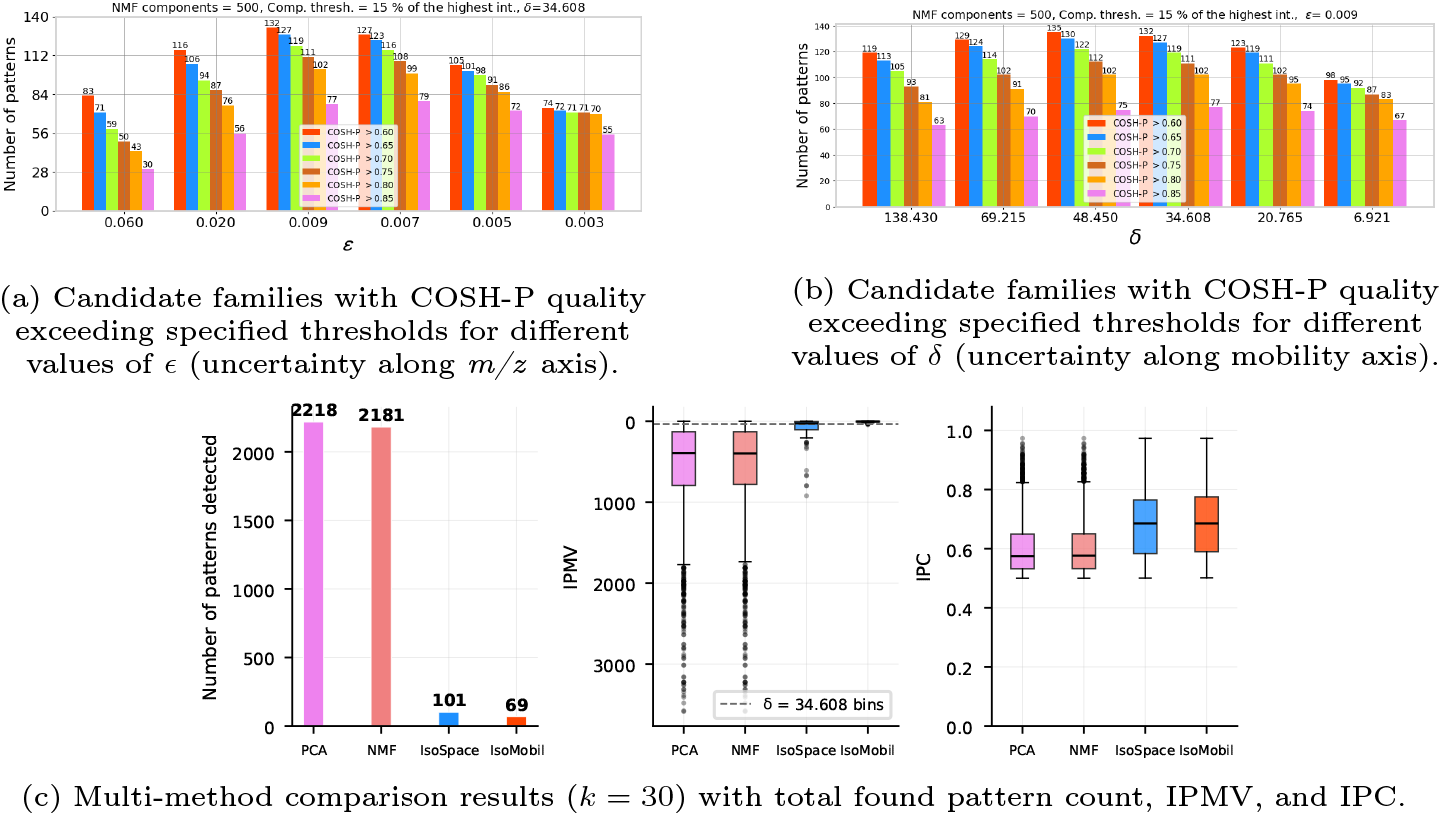
Case study 3 results-performance in human colon tissue. (a,b) Number and quality of found isotopologue candidate families as a function of *m/z*-tolerance *ϵ* and *mb*-tolerance *δ*: *ϵ* = 0.009 and *δ* ≈ 34.608 seem to yield best results. (c) Comparison of spatial-only (PCA, NMF), spatial+*m/z* (IsoSpace), and spatial+*m/z* +*mb*-filtering (IsoMobil). IsoMobil reduces found patterns’ IPMV while preserving their IPC.

**Fig. 5.**
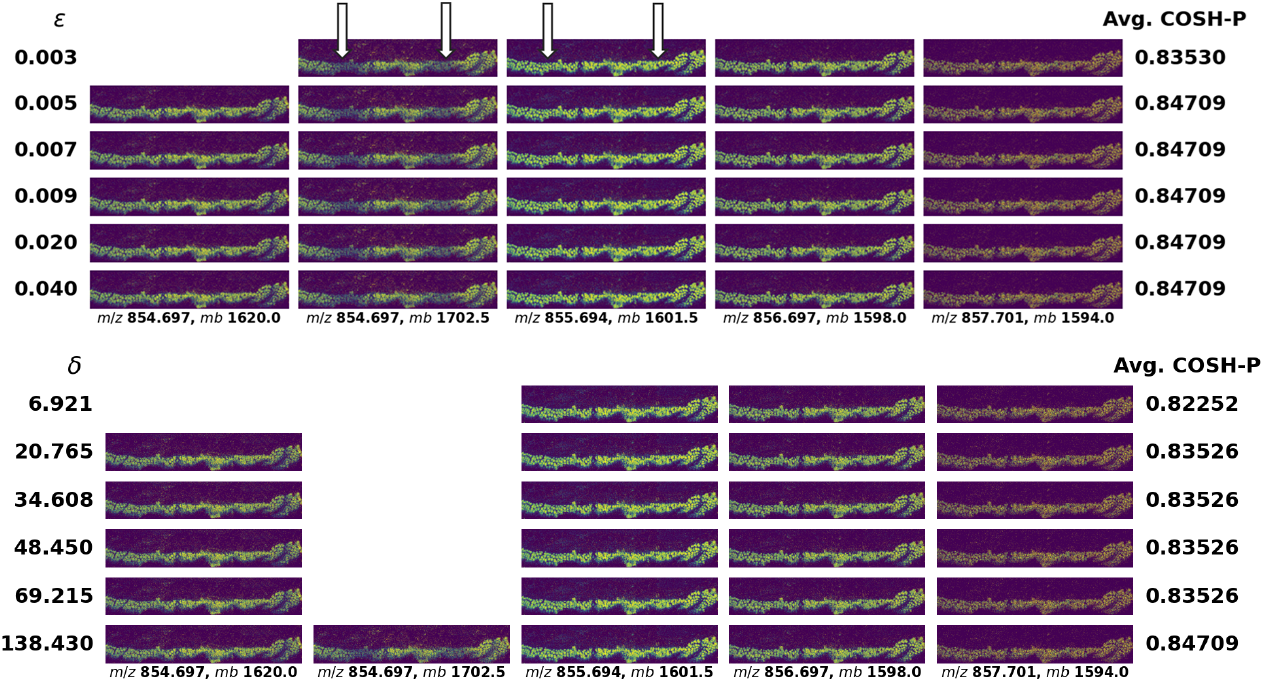
Case study 3 example-narrowing *mb*-tolerance removes false positive ion species stemming from isobaric molecular species. (top) Spatial+*m/z* filtering (relaxed *mb*-constraint) retains the ion species at *m/z* 854.697 and *mb*=1702.5 as part of this isotopologue candidate pattern despite narrowing *m/z*-tolerance *ϵ*. Subtle spatial variations (arrows) suggest this is incorrect. (bottom) Narrowing *mb*-tolerance *δ* removes the false positive, preserving mobility-consistent isotopologue candidates only.

## 4 Conclusion

IsoMobil is a novel approach to isotopologue discovery in complex IM-IMS datasets, grouping ion species that are consistent with reporting the same molecular species. This reduces the dataset’s dimensionality and provides concise lists of isotopologue candidates for independent confirmation through database matching, expert review, or MS/MS validation. IsoMobil is a unique isotopologue detector that enforces not only spatial and mass spectral consistency, but also ion mobility consistency, leading to fewer false positives. By avoiding assumptions about molecular formulas, chemical class, or isotope-intensity envelopes, it is one of the few methods that can be used in heterogeneous tissue environments where these assumptions may not hold. Its chemically informed dimensionality reduction avoids artifacts that traditional methods such as PCA can introduce (*e*.*g*., isotopologues from the same molecular species not being kept together). Overall, IsoMobil is an interpretability tool that makes downstream biological analysis of IM-IMS-based spatial omics datasets more robust by (i) simplifying complex spectra, (ii) reducing collinearity prior to clustering or classification, (iii) aiding species identification, and (iv) increasing confidence that features are genuine biological analytes since random noise does not form isotopologue patterns.

## Disclosure of Interests

The authors have no competing interests to declare that are relevant to the content of this article.

## Notes

### Competing Interest Statement

The authors have declared no competing interest.

